# Toxicity Profile of Artesunate in Rats and Dogs

**DOI:** 10.1101/2021.02.16.431559

**Authors:** Johanna S. Lang

**Affiliations:** Granzer Regulatory Consulting & Services, 81379 Munich, Germany

## Abstract

**OBJECTIVES:** The objectives of these studies were to investigate the toxicity, safety and toxicokinetics of single and multiple doses of artesunate for injection in rats and dogs.

**METHODS:** Sprague-Dawley rats and Beagle dogs were treated intravenously or intramuscularly for 28 consecutive days with doses of up to 30 mg/kg artesunate, evaluating toxicity, kinetics, genotoxicity, and cardiovascular and central nervous safety parameters after single and 4-week repeated administrations. Furthermore, respiratory parameters were evaluated after a single intravenous administration in rats.

**RESULTS:** Artesunate was well tolerated with no mortality and only minor effects on clinical pathology parameters. Following repeated intramuscular administration, local reactions at the injection site became evident.

**CONCLUSIONS:** The results obtained in these studies support the safe use of artesunate in humans.

Subchronic data for repeated intravenous and intramuscular artesunate in animals are available in the literature for a maximum duration of 14-days. Published results of genotoxicity studies *in vitro* in bacteria or cells of animal or human origin and *in vivo* in animals are equivocal, with a couple of negative results following oral administration and positive results in human cells.

With the studies performed in rats and dogs for a treatment period of 28-days including measures of safety pharmacology, toxicokinetic evaluation and *in vivo* genotoxicity data, a comparison of toxicity and exposure of intravenous versus intramuscular artesunate for a prolonged treatment period is possible, delivering important information, when a decision between both routes has to be made for use in patients. Results of the peripheral blood micronucleus assay in rats were positive, suggesting genotoxic potential for the intravenous route of administration of artesunate, what is, beside patients, likewise relevant for production and health care personnel.

## Introduction

Artesunate, an artemisinin-derivative, has been used for decades in humans for the treatment of severe *falciparum* malaria, which is a uniformly lethal disease if not treated promptly with potent antimalarial drugs. The WHO recommends the use of parenteral artesunate for the treatment of severe *falciparum* malaria since 2005.^1^ Although the mechanism of action remains to be completely elucidated, various data prove the efficacy of artesunate *in vitro*, *in vivo* in rodent and monkey challenge models, and in clinical and field isolates, by the number of successfully cured patients. Adsorption, distribution and elimination data are known for several routes of administration in commonly used laboratory species, including pregnant animals. Metabolism data are reported *in vitro* for animal and human cells, for animals as well as drug-drug interaction data for humans.

For intravenously administered artesunate, tolerability and efficacy were investigated in rats for three consecutive days. These studies, performed by Xie and Li, evaluated known haematotoxic and potential nephrotoxic effects of high doses of artesunate in uninfected and *plasmodium berghei-infected* rats. ^2,3^ Using the intramuscular route of administration in rats, studies of a duration of up to 7 days are available in the literature for artesunate as well as for the active metabolite dihydroartemisinin (DHA).^4,5^ For nonrodents, intravenous artesunate data for up to 14 days of treatment in dogs and in monkeys are reported.^6,7^

Extensive work was done evaluating reproductive and developmental effects *in vitro*, using whole embryo cultures, in rats, rabbits, and monkeys using the oral, intravenous, intramuscular or intraperitoneal route of administration.^7–26^ However, to date, no *in vivo* genotoxicity data for the intravenous route of administration are available in the literature.

During the last decade, nearly no additional work has been published to close the gap of nonclinical toxicity and safety pharmacology data for intravenous and intramuscular administered artesunate for a treatment duration beyond 14 days. In contrast, oral artesunate was investigated more recently for subchronic treatment (durations of one to three months) in mice, rats, rabbits and dogs.^27,28,29,30^

The aim of the studies presented in the following was to evaluate the toxicity of intravenous and intramuscular injected artesunate, in order to address potential risks by prolonged intravenous use, safety measures and for genotoxicity.

### Materials and methods

Two repeated dose toxicity studies and one respiratory safety study with administration of artesunate *via* the intravenous (IV) or intramuscular (IM) route were conducted in rats and dogs. Details of the study designs are shown in Table 1.

**Table 1:** Study Designs

| Study | Duration | Routes | Endpoints |
| --- | --- | --- | --- |
| Repeated dose toxicity, rat | 28-day treatment, 28-day recovery | IV and IM | General toxicity, neuro-behavioral changes (modified Irwin Test), body temperature, genotoxicity (micronucleus test) |
| Repeated dose toxicity, dog | 28-day treatment, 28-day recovery | IV and IM | General toxicity, safety pharmacology (cardiovascular effects), body temperature |
| WBP, rat | Single dose | IV | Respiratory system |

Abbreviations: IM = intramuscularly; IV = intravenously; WBP = whole-body plethysmography.

### Test article

Artesunate (artesunic acid, ZA1200207) was obtained from Guilin Pharmaceutical Co., Ltd. Purity of the batch used was ≥ 99%. Formulations of artesunate for intravenous or intramuscular administration were prepared using sodium bicarbonate as solvent (60 mg/mL) for the stock solution, and dilutions were made with sodium chloride to the final required concentrations for injections. Concentration and stability of the test item in the dosing formulations were analysed in representative samples from the animal studies by using an HPLC method, validated compliant with international guidelines and methods.

### Animal studies

Dose levels were selected based on a preliminary 5-day repeated dosing study in rats. For the repeated dose toxicity studies in rats and dogs, intravenous doses of 3, 10, 30 mg/kg/d and an intramuscular dose of 30 mg/kg/d were selected. Control animals received the vehicle (mixture of sodium bicarbonate and sodium chloride) by IV injection. The application volume was 5 mL/kg in rats and 2 mL/kg in dogs for intravenous and 0.5 mL/kg (stock solution 60 mg artesunate /L).

Purpose-bred conventional naïve Beagle dogs, 6-10 months of age (5-10 kg bw), and specific pathogen-free Sprague-Dawley rats, 6-9 weeks of age (males: 180-300 g bw; females: 160-200 g bw) were included in the studies. The animals were received from licensed providers, quarantined and acclimatised for at least 5 (rats) or 14 days (dogs) before inclusion into the studies. 21 dogs and 92 rats of either sex were randomly assigned (based on body weights) to the five groups (n=3 dogs/sex; n=10 rats/sex) by using a Provantis module for grouping. Additional animals (see Table 2) were kept for a recovery period of 4 weeks or used for TK blood samplings in rats. Animals were treated for 28 consecutive days intravenously into the tail vein (rats) or into peripheral veins of the limbs (dogs) or intramuscularly into the quadriceps femoris.

**Table 2:** Repeated Dose Toxicity Groups and Numbers of Animals

| Dose | Route | Rat (N per sex) |  |  | Dog (N per sex) |  |
| --- | --- | --- | --- | --- | --- | --- |
|  |  | Main | Recovery | TK | Main | Recovery |
| Control | IV | 10 | 5 | 3 | 3 | 2 |
| 3 mg/kg/d | IV | 10 | - | 6 | 3 | - |
| 10 mg/kg/d | IV | 10 | - | 6 | 3 | - |
| 30 mg/kg/d | IV | 10 | 5 | 6 | 3 | 2 |
| 30 mg/kg/d | IM | 10 | 5 | 6 | 3 | 2 |

Abbreviations: IM = intramuscular; IV = intravenous; N = number; TK = toxicokinetic.

The animals were group housed by sex in plastic solid-bottom cages (≤ 5 rats/cage) or in stainless steel cages (dogs), with single housing for collection of feces and urine samples, and for dogs additionally during designated procedures. Room temperature was kept between 18-26°C (dogs) or 20-26°C (rats) with a relative room humidity of between 40 and 70%, air changes > 8/h (dogs) or 15/h (rats), and a 12-hour light/dark cycle. Dogs were fed with approx. 250 g Dog Maintain Feed per day, and rats received SPF Rodent Maintenance Feed *ad libitum*. Animals were provided with purified water prepared in-house in the facility *ad libitum*. A pre-test health screening by a veterinarian was conducted for all animals used in the studies.

Animals (rats and dogs) were observed once daily for mortality and twice (during treatment) or once (during recovery) daily for clinical findings (study procedures are listed chronically in supplementary **Error! Reference source not found.**). Body weights were recorded once before dosing and once weekly thereafter, and body weight gains calculated. Food consumption was checked once daily in dogs and at least once weekly in rats, and average food consumption (g/animal/day) calculated for rats. All animals (rats and dogs) were subjected to ophthalmologic examinations prior to first dose and at the end of the treatment and recovery periods.

Blood for clinical pathology examinations was collected twice before the first dose in dogs, and on D28 or D29 and D56 (recovery period) in rats and dogs. For clinical chemistry, blood was collected into tubes with separation gel and coagulant, centrifuged to obtain serum and analyzed with a Hitachi-7180 Automatic Clinical Analyzer. Samples for haematology or coagulation analysis were collected into tubes containing EDTA-K2 or citrate sodium as anticoagulant and analyzed by the Sysmex XN-1000V or Sysmex CS-5100 Automated Analyzers. Urine and faeces (dogs only) were collected before dosing (dogs only) and once at the end of the dosing and recovery periods. Urine samples were analysed by the URIT-500B Urine Analyzer and dog faeces were analysed for occult blood in addition. Parameters determined are listed in supplementary **Error! Reference source not found.**.

Blood for determination of artesunate or its main metabolite dihydroartemisinin (DHA) was collected prior to dosing and 5, 30 min, 1, 3, 6 and 24 hours after dosing on D1. On D27 (dogs) or D28 (rats), samples were collected from all animals prior to dose and at 15, 30, 45 min, 1, 3, and 6 hours post dose. Blood was collected into pre-cooled heparinized tubes and stabilized by adding 40 mg/mL sodium fluoride (NaF) in water. Sample analysis was done using LC-MS/MS, validated compliant with international guidelines (ICH) and methods. Primary TK parameters, maximum plasma concentration (C_max_), time point of C_max_ (T_max_), and area under the concentration-time curve (AUC_(0-t)_), were calculated.

All rats and dogs scheduled for necropsy at the end of the treatment and recovery periods were anaesthetized (approx. 30-45 mg/kg pentobarbital sodium or ketamine in combination with xylazine [mixture of 20 mg/kg / 10 mg/kg for dogs; 90 mg/kg / 6 mg/kg for rats]), exsanguinated and subjected to macroscopic pathological examinations (see supplement **Error! Reference source not found.**), and organ weights analysis. Organs and tissues (see supplement **Error! Reference source not found.**) were taken for histopathological investigations (reported elsewhere). In addition, bone marrow samples were collected from the femurs of rats and from the ribs of dogs and smears were prepared for analysis.

### Safety pharmacology (cardio-vascular, neuro-behavioral and respiratory)

ECG parameters (see supplement **Error! Reference source not found.**) and body temperature were determined on D1 (1.5 h post dose), D2, D27 (prior to dose and 1.5 h post dose) and D28 as well as D56 (recovery period) by non-invasive telemetry in all dogs.

Rats were investigated using a modified Irwin Test (see supplement **Error! Reference source not found.**) to check for neuro-behavioural effects and measurements of body temperature 5, 30, 60, 240 minutes and 24 hours after the first and second to last dose.

A separate study to evaluate effects on respiratory parameters (see supplement **Error! Reference source not found.**) was conducted in Sprague-Dawley rats. Based on the 5-day preliminary toxicity study, dose levels of 10, 30 and 60 mg artesunate/kg bw were selected for single intravenous administration. Five animals per sex and group received a dose volume of 5 mL/kg into the tail vein. Control animals received the vehicle. Body weights were recorded on the day of dosing and respiratory function measurements (see supplementary **Error! Reference source not found.**) were evaluated by whole-body plethysmography with data acquired immediately for baseline (90 minutes before dosing). Post-dosing data were generated at 10 min, 0.5, 1, 4 and 24 hours after dosing. Animals were stratified to five evaluation blocks, with at least one animal of each group represented in each block of measurement.

### Genotoxicity

At the end of the dosing period (D29), the first five rats/sex/group were selected from scheduled necropsy animals for micronucleus analysis using the MicroFlow^®^ PLUS Rat Blood Micronucleus Analysis Kit (Litron Laboratories, USA). In peripheral blood, approximately 20,000 CD71 high-positive reticulocytes (RET) were counted for each animal to determine the number of micronucleated reticulocytes (MN-RET) and frequencies (%MN-RET). As a measurement of cytotoxicity, the percent of reticulocytes (%RET) were calculated from total erythrocytes. Negative and positive controls were included and mean and standard deviation of %RET and %MN-RET calculated.

### Data and statistical analysis

Statistical comparison was performed between treatment groups and control groups. Group means and standard deviations were calculated, separately for each sex using the statistical methods described in supplementary **Error! Reference source not found.**. The data of mortality, clinical observations, food consumption, ophthalmology, urinalysis, occult blood in feces (dogs only) and bone marrow smears were not statistically analyzed.

### Ethics and compliance statements

All animal work in this study was conducted complying with the “Guide for Care and Use of Laboratory Animals” issued in 2011 by the National Research Council, USA and protocols, amendment(s) and procedures reviewed and approved by the Institutional Animal Care and Use Committee (IACUC). The studies were conducted according to OECD GLP regulations in a GLP certified laboratory and in consideration of internationally accepted testing guidelines (OECD, ICH) for preclinical safety studies.

## Results

### Formulation analysis

Results of the validation study demonstrated good specificity of the HPLC method, and DHA did not interfere with the detection of artesunate. The calibration curves of artesunate showed good linearity in the range of 10~80 μg/mL, acceptance criteria for sensitivity, carryover, intra-accuracy, precision and homogeneity were met. The dose formulations of artesunate and postprocessed samples were stable during the period and conditions used.

Representative samples for formulation analysis were taken in all three studies on the first, second and last day of formulation. The results of system suitability tests, stock solution comparisons, performance checks and standard curves etc. met the requirement of the analytical method. No artesunate was detected in the control dose formulation of all three studies. Measured concentrations of the dose formulation samples of the first preparation in the rat study were out of specification (88.5, 88.4, 87.4 or 82.1%, respectively of the nominal concentrations) in the low, mid and high dose IV and the undiluted stock for IM preparations, including backup samples used for reanalysis. The mean measured concentrations of artesunate dose formulations from the second preparation, as well as all other samples determined for the rat and dog studies were within 90.0%-110.0% of the nominal concentrations, with a relative deviation of no more than 2.0%, and thus within the acceptance criteria.

### *In vivo* examinations

None of the animals died before scheduled termination. Clinical signs indicative of general toxicity were not observed in rats and dogs. In rats, swelling at the injection sites was observed (first time observed on D10) in all male and female animals of the group receiving intramuscular injections (30 mg/kg/d). Swelling of the hind limb (left and/or right) was seen in 8 of 15 male and 10 of 15 female rats (accompanied by limping in 4 females) and in all dogs during the second half of the treatment period. All swellings recovered during the treatment-free period. One male dog showed additionally decreased activity (D12-28), which was associated with an abscess and injury on the right hind limb and limping.

In rats, mean body weights and body weight gains were not affected in male and female animals receiving intravenous doses of 3, 10 and 30 mg/kg/d in comparison to control animals, while intramuscular treated animals gained less body weight (males statistically significant D7 to D35, females below 10%), caused by a significantly reduced food consumption (males whole treatment period, females D17-21). The food consumption and body weight (partly in males) completely recovered.

In dogs, body weights were slightly (below 5% difference) lower in female animals of the high dose IV and IM treatment groups (30 mg/kg/d) compared to the other treatment groups and the control group. In the female IM group, occasionally food was left over. Body weights recovered in females of the IM group, but remained slightly lower in females of the high dose IV group.

No ophthalmological effects were observed in rats and dogs at any time point. Following single and repeated dosing, body temperature was not affected in both species.

### Clinical pathology

In rats, haematological changes affected the animals of the 10 and 30 mg/kg/d IV group and of the 30 mg/kg/d IM group. Red blood cell parameters indicative of anaemia (red blood cell count RBC, hemoglobin HGB and hematocrit HCT) were about 50% decreased with a concomitant 2-4-fold increase of the reticulocytes (RET). The interrelated parameters (MCV, MCH, MCHC and RDW) were also affected. In addition, the number of platelets (PLT) and the platelet Crit (PCT) were up to 50% increased in these groups. The number of WBCs was increased reaching statistically significance (10 mg/kg/d only males), by an increase of the absolute and relative neutrophiles (NEUT, 2-4-fold), lymphocytes (LYMPH, up to 2-fold), monocytes (MONO) and basophiles (BASO, both up to 10-fold). The thrombin time (TT) and the activated partial thromboplastin time (APTT) but not the prothrombin time (PT) were statistically significantly shortened. Fibrinogen was statistically significantly increased in the IM (male and female) treatment group. The absolute and relative numbers of nucleated red blood cells were up to 200-fold increased in a dose-related manner in all artesunate groups. Apart, in the low dose group treated with 3 mg/kg/d IV, only a few of the interrelated parameters were affected (see supplementary **Error! Reference source not found.**). Except the RDW in males and the MCV and MCH in females of the high dose IV and IM groups, all changes recovered.

Clinical chemistry examination revealed a statistically significant decrease of cholesterol (CHOL) levels in male rats in the mid and high dose IV groups (10 and 30 mg/kg/d) and the IM group (30 mg/kg/d), while triglyceride (TG, excluding females of the 10 mg/kg/d group), total bilirubin (T-BIL) and direct bilirubin (D-BIL, bilirubin including females of the 3 mg/kg/d group) values were increased (statistically significant in the IM group). Total bile acids (TBA) and phosphate (P) were increased in male and female animals of the mid (only P) and high dose IV and IM groups, reaching statistical significance only in males. The level of albumin (ALB) was decreased in animals (statistically significantly in females) of the IM group with statistically significant elevated globulin (GLO) values and a decreased ratio of A/G in male and female rats. Additional statistically significant findings in the treatment groups compared to controls appear to be incidental and not related to treatment with artesunate. For details please see supplementary **Error! Reference source not found.**. Except P and TBA (still increased in males of the IM group), all parameters completely recovered.

As observed in rats, haematological parameters indicative of anaemia were affected in dogs. The values of RBC, HGB, HCT were ~30-40% decreased in animals of the IV (10, 30 mg/kg/d) and IM (30 mg/kg/d) treatment groups. MCH, MCHC, PLT, and PDW were affected only slightly and not in all groups. The effect on RET was not as clear cut as in rats, with 2-3-fold increased values in the low and mid dose groups and ~50-70% decreased levels in the high dose IV and IM treatment groups. However, MPV was increased in the mid and high dose IV and the IM groups. For details please see supplementary **Error! Reference source not found.**. The effects on white blood cell parameters did not indicate a clear effect, also pre-treatment WBCs showed quite some variability. The NEUTs were minimally increased, the absolute and relative MONOs about 3-fold increased, and EOSs minimally decreased. As observed in rats, the absolute and relative numbers of nucleated red blood cells (NRBCs) were up to 300-fold increased in all artesunate groups. In contrast to rats, coagulation parameters (PT, APTT and TT) were not affected by the treatment in dogs. The only finding observed was an increase (partially statistically significant) of the fibrinogen values in the high dose IV and IM groups. Except fibrinogen levels (lower in males, but higher in females of the 30 mg/kg/d IV and IM groups) all hematological parameters recovered back to control or baseline values.

Aspartate aminotransferase (AST), alkaline phosphatase (ALP except females of the IM group), LDH (only females) and CK (excluding males of the high dose IM group) were increased in the 30 mg/kg/d IV and IM groups. T-BIL values were statistically significantly increased and above the range of pre-dose level in the 30 mg/kg/d IV group. For details please see supplementary **Error! Reference source not found.**. Further affected parameters were creatinine and potassium levels (statistically significantly decreased only in male dogs of the IM group), and decreased calcium levels in females in the 10 or 30 mg/kg/d IV group and the IM group. ALB was decreased, and globulin increased with affected A/G ratios in males of the IM treatment group (the same as observed in the rats), while this effect was seen in the high dose IV males to a lower degree. A trend towards lower ALB and higher GLO as well as lower A/G ratios was seen in females of these dose groups (30 mg/kg/d IV or IM). Females of the IM treatment group showed also a statistically significant increase (above control and predose ranges) of amylase. A complete recovery was observed for all the affected parameters, except ALP and CK (still higher in males of the IM group), and CREA (higher in males of the 30 mg/kg/d IV and IM groups).

No treatment-related findings were observed in urinalysis in male and female rats and dogs. No occult blood was determined in the faeces of dogs.

### Post-mortem evaluations

Terminal body weights at necropsy were lower in the IM treatment group (30 mg/kg/d). Liver weights were increased in the 10 and 30 mg/kg/d IV groups and the IM group (absolute or relative, statistically significant in the 30 mg/kg/d groups), and marginally in both sexes of the low dose group (3 mg/kg/d). Absolute and relative spleen weights were statistically significantly increased in the mid and high dose IV and the IM groups. Heart weights (absolute and relative) were increased in the high dose IV group. In addition, a few statistically significant differences were observed: decreased absolute and relative thymus weights and increased relative testes weights in males of the IM group. For details see supplementary **Error! Reference source not found.**. Increased spleen weights (both sexes), in females of the IM group increased liver weights and in males of the IM group decreased kidney weights showed no or incomplete recovery.

At the end of the treatment period, no macroscopic observations were noted in male and female rats of the control and low dose IV groups (3 mg/kg/d). Macroscopic findings of enlarged spleen were observed in in 3/10 males in the 10 mg/kg/d IV group, 7/10 males and 5/10 females in the 30 mg/kg/d IV group and 7/10 males and 6/10 females in the 30 mg/kg/d IM dose group. In addition, enlarged administration sites (5/10 males) and crusts (1/10 males) as well as red discoloration at the administration site (1/10 females) were observed in rats of the IM dose group. At the end of the recovery period, no macroscopic findings were observed in any of the animals.

In dogs, terminal body weights at necropsy were lower without dose-relation in females of all dose groups. Absolute and relative spleen and liver (males only) weights were dose-dependently increased in the 10 and 30 mg/kg/d IV groups and the IM group. Absolute heart weights were not affected in male animals, while they were dose-dependently lower in female animals of all dose groups (including relative weights). For details see supplementary **Error! Reference source not found.**. No or incomplete recovery was observed for liver and spleen weights (still higher in male animals of the high dose IV and IM groups). In female animals, heart and liver weights were higher in the high dose IV group and spleen weights were still higher in the IM group.

Macroscopic examination of dogs at the end of the treatment period did not reveal any findings in animals of the IV treatment groups (3, 10, 30 mg/kg/d). All three male and female animals of the IM group (30 mg/kg/d) showed enlargements at the administration sites, and one male animal showed injury at the administration site in addition. No further macroscopic findings were observed. At the end of the recovery period, no macroscopic findings were observed in any of the animals.

Bone marrow analysis revealed no treatment-related findings in rats and dogs of the control and low dose IV group (3 mg/kg/d). In all rats and dogs of the mid and high dose IV (10 or 30 mg/kg/d) and the IM group (30 mg/kg/d) groups, erythroid dysplasia was noted and the granulocyte to erythroid ratio was inverted (GE ratio <1: 1) characterized by active proliferation of bone marrow, and/or increased segmentation cells (rats only) and erythron, and/or relatively decreased granulocytes and lymphocytes. In dogs, hemophagocytic cells were observed in most of the smears (16/18 cases). Furthermore, increased erythroblastic islets were noted in 4 of 20 rats of the IM dose group, but not in the IV dose group animals. In addition, for one dog of the IM group, the GE ratio was 12.14:1 with significantly decreased erythroid hyperplasia, increased granulocytes (mainly neutrophile granulocytes), high proportions of naked nuclei in megakaryocytes, and a trend of stagnated hematopoietic function. In another dog of this group, the quality of smear was poor and the smear was thick; the proportion of naïve granulocytes in granulocytes was increased. In two dogs of the high dose IV group (30 mg/kg/d), and single rats of the large 10 mg/kg/d IV group and the IM group, smudge cells were observed.

### Toxicokinetics

All samples for the control group in rats and dogs on the first day of dosing (D1) and at the end of the treatment period (D28 rat, D27 dog) were below the quantitative lower limit of detection (10.00 ng/mL). The TK results for rats and dogs are presented in detail in supplement **Error! Reference source not found.** and **Error! Reference source not found.**.

As expected after intravenous injection of artesunate, C_max_ was obtained at the first sampling time point 5 minutes after dosing in rats and dogs. Artesunate was rapidly metabolised to DHA within minutes after intravenous injection and C_max_ was also observed at the first sampling time point of 5 minutes in all IV dose groups (3, 10, 30 mg/kg/d), except in the high dose IV group (30 mg/kg/d) at the end of the treatment period, where C_max_ was determined 15 minutes after dosing. After IM injection of artesunate (30 mg/kg/d), C_max_ levels were reached within 5 (rats) and up to 15 (dogs) minutes after dosing, but metabolism to DHA was somewhat slower with T_max_ levels achieved within the first 30 minutes after dosing.

In rats, the increase of artesunate C_max_ was almost dose-proportional between the 3 and 10 mg/kg/d groups (range 2.9-4.9), and higher than dose proportional between the 10 and 30 mg/kg/d IV groups (range 5.9-14). With respect to DHA, the increase of the metabolite levels in the IV dose groups reflect the increase of doses in an almost proportional way. Animals of the IM dose group treated with 30 mg/kg/d showed lower C_max_ levels than in the high dose IV group (30 mg/kg/d). With the exception of D1 values in females (15%), plasma levels of males and females were about 62-69% lower in the IM group. This is also reflected in the levels of DHA which were about 55-78% lower.

Although the reanalysis of artesunate dosing solutions on the first day of dosing (D1) in rats was only about 82-88% of nominal, this was not reflected in the C_max_ levels of female animals of all dose groups and routes and in the male 30 mg/kg/d IV and IM groups. All C_max_ levels were lower after 28 days of treatment with artesunate, except in low and mid dose IV male rats (3 or 10 mg/kg/d) which were higher than the D1 C_max_ values.

For artesunate on D1 in rats, AUCs could only be calculated for the IM dose group (30 mg/kg/d) because most of the data at the various time points were below the limit of detection for the IV dose groups due to the fast metabolism of artesunate to DHA. In general, AUCs for artesunate and DHA were lower than D1 levels after 28 days of treatment with 30 mg/kg/d IM (see Figure 1). In the IV dose groups, AUCs increased with dose, and lower values were seen in female animals in the low and mid dose IV groups (3, 10 mg/kg/d) compared to male animals, while there was no difference between male and female animals of the high dose IV and IM dose groups (30 mg/kg/d). In animals of all treatment groups, except males of the low dose group (3 mg/kg/d), the AUCs for DHA were somewhat lower on D28 than at the start of dosing on D1.

**Figure 1:**
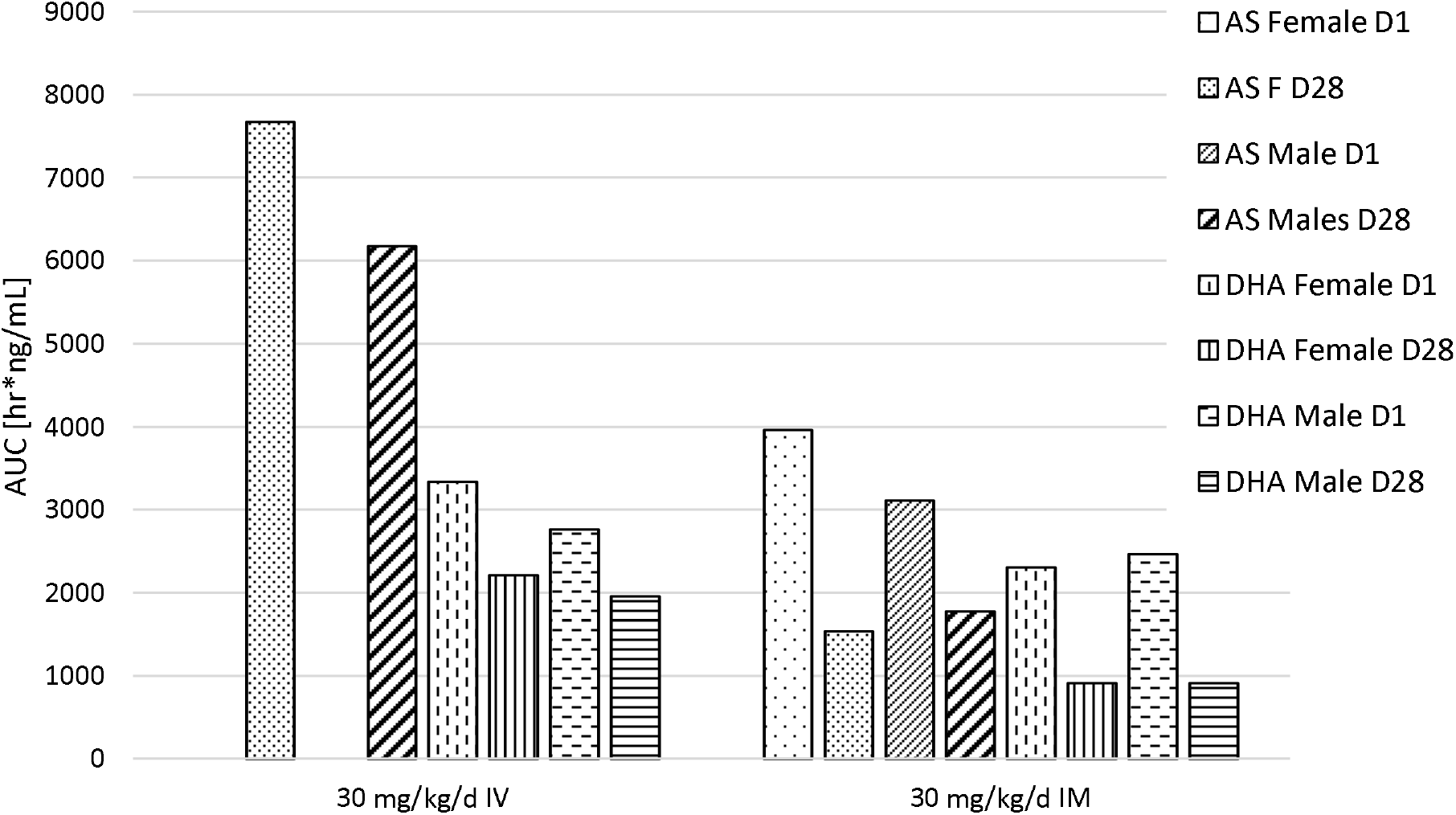
Comparison of plasma exposure in rats. Exposure in terms of AUC is displayed for female and male rats after a single (D1) and repeated (D28) intravenous (IV) or intramuscular (IM) treatment with 30 mg artesunate/kg bw/day for artesunate (AS, left four bars per dose group) and its main metabolite dihydroartemisinin (DHA, right four bars per dose group). No AUC was calculated for males and females on the first day of treatment (D1) for artesunate due to fast metabolism to DHA. For the last sampling, additional early sampling time points were included, to enable calculating AUC values.

In dogs, plasma levels of artesunate (C_max_ and AUCs) were higher than in rats. There was no clear difference between D1 and D27 for the IV groups (3, 10, 30 mg/kg/d), but the D27 plasma levels were lower in the IM dose groups (30 mg/kg/d) as compared to D1. C_max_ and AUC increased slightly higher than dose proportional (4.1-9.2) with a slightly lower increase between the low and mid dose IV females on D27 (C_max_: 2.4, AUC 2.1). As observed in rats, plasma levels in dogs (C_max_ and AUC) were lower in the IM group treated with 30 mg/kg/d than in the corresponding high dose IV group. C_max_ values were in the range of 64 to 92% and AUCs in the range of 43 to 74%.

The C_max_ levels of the metabolite DHA in dogs were almost in the same range as in rats with the exception of the 30 mg/kg/d IV group, for which the levels were lower in dogs than in rats. The increase in C_max_ and AUC was either dose proportional in females on D27 or less than dose proportional (range 2.1-2.7, 2.5-3.8, respectively) in most cases. The IM C_max_ level of DHA in animals treated with 30 mg/kg/d in comparison to the corresponding high dose IV group were about 29-68% lower. However, AUCs were higher in the IM dose group compared to IV in females on D1 and D27 and males on D1 (range 3-12%) and lower in males on D27 (16%, see Figure 2).

**Figure 2:**
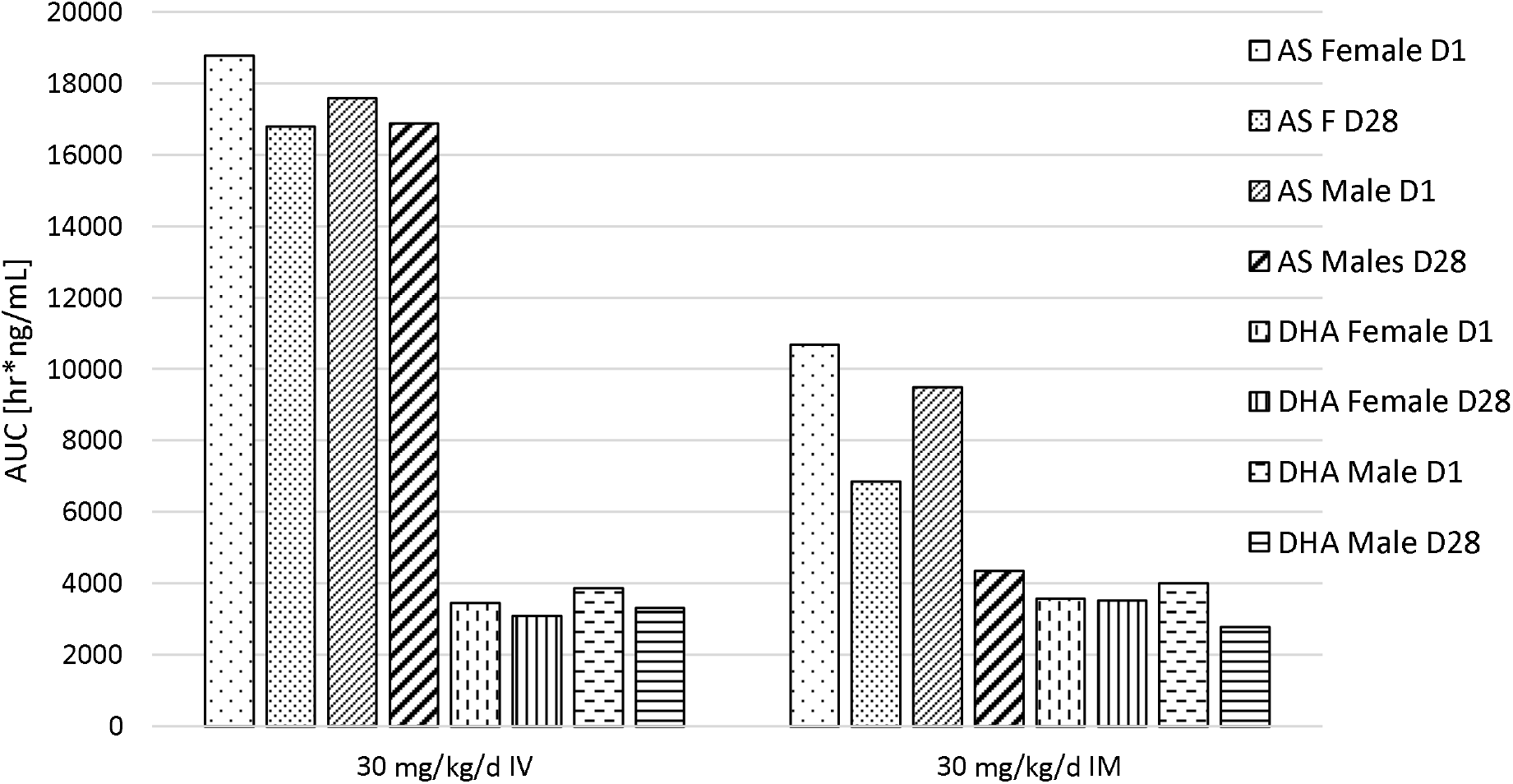
Comparison of plasma exposure in dogs. Exposure in terms of AUC is displayed for female and male dogs after a single (D1) and repeated (D27) intravenous (IV) or intramuscular (IM) treatment with 30 mg artesunate/kg bw/day for artesunate (AS, left four bars per dose group) and its main metabolite dihydroartemisinin (DHA, right four bars per dose group).

Results did not indicate an obvious gender differences between artesunate and DHA levels in male and female rats and male and female dogs.

### Safety pharmacology

Cardio-vascular safety was investigated in male and female dogs on D1 (1.5 h post dose) and D2 and on D27 (prior to dose and 1 h post dose) and D28 and D56 (recovery period). No treatment or test item-related effect were observed in animals in the IV or IM treatment groups in comparison to the control group. PR and RR intervals, wave duration, QRS duration, and QT interval, heart rate and QTcv were within normal limits for all groups and no pathologic arrythmias were noted at any of the time points. One individual statistically significant increase of the corrected QT interval (9.4%) observed in female animals of the high dose IM group (30 mg/kg/d) on D28 of the study period can be regraded as incidental because this value was in the range of the values observed on D27 predose and 1 hour post-dose.

Neuro-behavioural assessments were conducted in male and female rats pre-dose and at several time points (5, 30, 60 and 240 minutes and 24 hours) after the first (D1/2) and second to last dose (D27/28) by applying a modified Irwin screen. No treatment or test item-related findings were observed in animals of the treatment groups in comparison to control animals.

In a separate study, respiratory safety was investigated by whole-body plethysmography in male and female rats treated with a single intravenous (10, 30 or 60 mg/kg) artesunate injection prior to dosing and at several intervals post-dose (10 min, 0.5, 1, 4 and 24 hours). The results did not indicate a test item or treatment-related effect on the tidal volume, minute volume and respiratory rates. 10 minutes following administration, respiratory parameters were increased in all animals independent of the treatment, some becoming statistically significant compared to their respective predose values. All values returned back to baseline 0.5 to 1 hour post-dose and were therefore assumed to be related to the intravenous dosing procedure rather than to treatment with the test item.

### Genotoxicity

The genotoxic potential of the test item was investigated in male and female rats. Blood samples for a micronucleus assay in peripheral erythrocytes were collected at the end of the treatment period on the day of necropsy (D29). The mean percentage of reticulocytes (%RET) was increased in the mid and high dose IV and the IM treated animals (13.6%, 7.1%, 5.8% for males, 6.4%, 9.9%, 9.2% for females, respectively) compared to 0.7% and 1.6% in males and 1.3% and 1.9% in females of the control and low dose IV group. The percentage of micronucleated reticulocytes (%MN-RET) was 0.54%, 3.34%, 2.99% in males and 0.64%, 0.66%, 0.56% in females of the mid and high dose IV and the IM treated animals (p<0.01 for the IV groups, p<0.05 for the male IM group), respectively compared to 0.16%, 0.18% (males) and 0.11%, 0.13% (females) %MN-RET in the control and low dose IV group, respectively (being in the range of historical and assay negative control data). As the average values for micronucleated reticulocytes exceeded the upper of the historical ranges of negative/solvent controls and reached values of the assay internal positive control, artesunate was classified positive for the *in vivo* peripheral blood micronucleus assay.

## Discussion

Artesunate has been shown to exhibit relatively minor effects at high doses on the central nervous, cardiovascular and respiratory systems. The dose levels of artesunate used in the presented studies (up to 30 mg/kg/d intravenously or intramuscularly for 4 weeks or a single intravenous dose of 10, 30 or 60 mg/kg) were below doses known to cause central nervous or breathing effects. Following intravenous administration of artesunate, doses of 250 mg/kg and above caused decreased activity, analgesia, muscle relaxation, and hypnosis in mice.^31^ Intramuscular administration of 150 mg/kg resulted in neurologic effects and mortality.^32^ In rats, rabbits, guinea pigs, dogs and monkeys, intravenous doses reported lowering body temperature, causing EEG abnormalities, convulsions and breathing arrest between 160 to 640 mg/kg.^31^ A no effect level (NOEL) was determined for 80 mg/kg intravenously in monkeys and dogs. In our studies, no effects on behaviour or cardiovascular effects were noted for 4-week repeated intravenous or intramuscular injections. As well, no breathing abnormalities following a single intravenous dose of up to 60 mg/kg artesunate were determined.

After repeated oral doses of artesunate, the levels of exposure decreased in rats and rabbits, this was especially evident in the high doses.^22,33^ A decline in exposure in terms of C_max_ and AUC was determined as well in our studies following intravenous (only in rats) and intramuscular administration of 30 mg/kg/d for artesunate and for its main metabolite DHA. Reduced plasma protein binding (normally between 73-82% in rats) was demonstrated at higher doses (66% at DHA concentrations > 125 ng/mL).^34^ Furthermore, the binding capacity of artesunate was significantly different between male and female rats in the study by Li et al., which could explain the differences observed between the sexes especially in the 3 mg/kg/d group and the supra-proportional increases in exposure in the 30 mg/kg/d IV group in rats.

A bioavailability of intramuscular administered artesunate is reported with 85-105% for artesunate and DHA.^35^ In our studies, bioavailability of artesunate based on AUC ranged between 26 to 57% in rats and between 20 and 29% in dogs, although AUC values are of limited value, as they were based on single data due to the fast metabolism of artesunate to DHA. Bioavailability of DHA was 69% and 89% in females and males after the first treatment and only 41% and 47% at the end of the 4-week treatment period. In dogs, bioavailability of DHA following intramuscular administration ranged between 84-114% after a single and repeated injections, thus being in the range determined by Li et al.

Comparing human and animal exposure and the resulting effects following intravenous artesunate treatment, plasma concentration is not considered to be a good indicator of tissue exposure (measured by biologic effects, e.g. reticulocyte reductions) for artesunate across species.^26^ In our studies, reticulocyte levels were reduced for absolute counts in dogs only at the end of the 4-week treatment period, but increased in absolute and relative counts in rats. Due to the mechanism of action, (affecting the blood stages of *plasmodium*, i.e. red blood cells), thus leading to anaemia (as demonstrated by our results in both rats and dogs), the increased reticulocyte levels determined in rats are indicative of regenerative anaemia by increased reticulocyte maturation. The inversed granulocyte to erythroid ratio (GE ratio <1:1) might be the result of an increased release of granulocytes, as indicated by increased plasma leukocyte counts. Increased spleen and liver weights, especially in rats but also noted in dogs, are indicative of extramedullary haematopoiesis to compensate anaemia, complying with results of other groups, e.g. as seen in a subchronic oral toxicity study for artesunate in dogs and a short-term intramuscular study in mice.^30,36^

Effects on clinical chemical parameters in rats and dogs were mainly characterized by reduced albumin levels, and increased globulin and bilirubin levels. The bilirubin is associated with an increased breakdown of red blood cells. In addition, an increase in liver parameters (AST, ALT) was seen in dogs, which was partly still evident at the end of the recovery period.

The effects described in our studies above were all of a minor toxic nature and most of them demonstrating a nearly to complete recovery, demonstrating a good tolerability and safety profile for artesunate. Comparing the intravenous to intramuscular route of administration, despite differing systemic exposure, the effects on target organs were nearly comparable to even more exaggerated in the intramuscular groups, demonstrating a good efficacy of this route, despite the local effects, but only arising after one week of repeated daily treatment. A good tolerability is known for intravenous or intramuscular use of artesunate in humans as well. In openlabel randomised controlled trials in patients admitted to hospital with severe *falciparum* malaria in Bangladesh, India, Indonesia, and Myanmar (performed by the South East Asia Quinine Artesunate Malaria Trial, SEAQUAMAT group) and in paediatric patients admitted to hospital in the sub-Saharan Africa (AQUAMAT trial), superior efficacy, and a significantly reduced mortality were shown over quinine.^37,38^ A better tolerability compared to intramuscular artemether has been demonstrated in several studies. Following administration of 2.4 mg/kg artesunate at 0, 12 and 24 hours intravenously continued once daily until oral tolerance in adults (maximum 7 days) demonstrated a good tolerability. Beside a rare type-1 hypersensitivity reaction in about 1 in 3,000 treated patients, no serious adverse effects were noted. Our present studies demonstrated that artesunate, applied over a period of four weeks, was safe and produced only minor and to a great extent reversible effects in rats and dogs. Despite varying plasma levels comparing the intravenous to the intramuscular route of administration, effects on target organs were to the same extent, suggesting a comparable exposure, and potential effectivity, of artesunate for both routes, and despite some local effects during extended (beyond one week) duration of treatment for the intramuscular route, a comparable safety profile.

## Acknowledgments

The experimental studies were funded by Guilin Pharmaceutical Co., Ltd.

